# Auxin-mediated rapid degradation of selective proteins in hippocampal neurons

**DOI:** 10.1101/612119

**Authors:** Risako Nakano, Naoki Ihara, Shota Morikawa, Ai Nakashima, Masato T. Kanemaki, Yuji Ikegaya, Haruki Takeuchi

## Abstract

Genetic manipulation of protein levels is a promising approach to identify the function of a specific protein in living organisms. Previous studies demonstrated that the auxin-inducible degron (AID) strategy provides rapid and reversible degradation of various proteins in fungi and mammalian mitotic cells. In this study, we employed this technology to postmitotic neurons to address whether the AID system could be applied to the nervous system. Using adeno-associated viruses, we simultaneously introduced EGFP fused with an AID tag, and an F-box family protein, TIR1 from *Oryza sativa* (OsTIR1) into hippocampal neurons. In dissociated hippocampal neurons, EGFP fluorescence signals rapidly decreased when adding a plant hormone, auxin. Further, auxin-induced EGFP degradation was also observed in hippocampal acute slices. Taken together, these results open the door for neuroscientists to manipulate protein expression levels by the AID-system in a temporally-controlled manner.

## Introduction

Conditional manipulation of the protein expression level is indispensable for understanding not only the function of specific proteins, but also complex biological systems. Various methods have been developed to regulate the expression level of specific proteins at the level of transcription or translation [1–3]. General approaches to control the protein level are the disruption of DNA sequence coding for a specific protein by gene editing and suppression of mRNA level by RNA interference [4–7]. However, as these methods deplete proteins in an indirect way, their temporal specificities heavily rely on the stability of the target proteins.

To achieve precise temporal control of protein expression, a variety of systems have been invented that target posttranslational protein degradation using cell-permeable small molecules [8–13]. The auxin-inducible degron (AID) system has been developed by modifying a plant-specific ubiquitin-proteasome pathway [14]. Skp1-Cullin-F-Box protein (SCF) complex catalyzes polyubiquitination of proteins destined for degradation by the 26S proteasome. The SCF complex is composed of the core CUL1-RING complex and an F-box substrate-recognition subunit. Although the SCF complex is highly conserved across species, a type of F-Box protein determines the specificity of substrate recruited to the SCF complex. TIR1 is an F-box protein that is only preserved within plant species and recognizes a degron sequence (hereafter called the AID tag) conserved in the AUX/IAA family proteins only in the presence of phytohormone auxin for degradation via the ubiquitin-proteasome pathway [15–17]. The AID system has been developed by introducing two components, the plant-specific TIR1 and the AID tag, the latter of which is fused with a protein of interest to promote the degradation of AID-tagged proteins through the binding of TIR1. To date, the AID system has been employed for conditional ablation of specific proteins in a variety of cells [14, 18–20]. Previous studies demonstrated that the AID system achieves not only conditional, but also rapid degradation of target proteins on a time scale of several ten minutes [14, 18, 21]. In this study, we examined whether the AID system can be used for manipulating specific protein levels in the nervous system.

## Materials and methods

### Animals

All experimental procedures were performed with the approval of the animal experiment ethics committee of the University of Tokyo (approval number: P29-15, and in accordance with the guidelines for the care and use of laboratory animals of the University of Tokyo. C57BL/6J mice were purchased from Japan SLC (Shizuoka, Japan).

### DNA construction and AAV vector production

Adeno-associated virus (AAV) vector was generated as described previously [22]. pAAV-hSyn-OsTIR1-P2A-mAID-EGFP construct was synthesized by replacing cording region of the pAAV-hSyn-EGFP (Addgene plasmid #50465) with the OsTIR1-P2A-mAID-EGFP-NES sequence OsTIR1-p2A-mAID-EGFP-NES sequence was amplified by PCR from pAY8 using following primers; 5’ GGATCCGCCACCATGACATACTTTCCTGAAGAGGTCGTC 3’ and 5’ GCTTTGTACGGAATTGGGAGGTGTGGGAGGAGGTTTT 3’. The PCR product was subcloned into the NcoI and EcoRI sites of the pAAV-hSyn-EGFP. HEK293T cells were transfected with the pAAV-hSyn-OsTIR1-P2A-mAID-EGFP and two AAV helper plasmids encoding serotype DJ (Cell Biolabs, San Diego, CA, USA) using polyethylenimine (Polysciences, Warrington, PA, USA). Three days after transfection, AAVs were collected from HEK293T cells and purified using AAVpro Purification Kit (Takara Bio, Shiga, Japan) according to the manufacturer’s protocol. The AAV titer was determined to be 4.8×10^13^ vg/ml by real-time PCR using ITR2 primers [23].

### Primary culture of hippocampal neuronal cells

Dissociated hippocampal neurons were prepared from postnatal day (PD) 0 C57BL/6J mice as previously described [24]. Mice were anesthetized by hypothermia and quickly decapitated. Hippocampal tissue was dissected and minced in pre-warmed Hank’s Balanced Salt Solution (HBSS) and treated with 0.25% Trypsin/EDTA at 37 °C. After 15 min of incubation, the tissue was treated with DNaseI (Sigma-Aldrich, St Louis, Missouri, USA) at room temperature (RT) for 5 min and washed with HBSS three times. HBSS was replaced with Neurobasal plating medium [Neurobasal Medium containing B27 Supplement (1:50), 0.5 mM Glutamine Solution, 25 μM Glutamate, Penicillin/Streptomycin (1:200), 1 mM HEPES, 10% horse serum (heat-inactivated and filter-sterilized, Gibco, Inc., Grand Island, NY, USA)]. Tissue was triturated using fire-polished Pasteur pipettes and filtered through a 40-μm-pore cell strainer (Corning, New York City, NY, USA). Hippocampal cells were plated on poly-D-Lysine coated glass base dishes (35 mmf with a window of 12 mmf, 0.15 mm thick glass; IWAKI) at a density of 8.0×10^4^ cells/well, and placed in a 37 °C, 5% CO_2_ incubator. At 1 day *in vitro* (DIV), Neurobasal plating medium was replaced with Neurobasal feeding medium (Neurobasal Medium containing B27 Supplement (1:50), 0.5 mM Glutamate Solution, Penicillin/Streptomycin (1:200), 1 mM HEPES). At 2 DIV, the medium was replaced with fresh feeding medium containing a final concentration of 5 μM cytosine β-D-arabinofuranoside (AraC; Sigma-Aldrich) for 24 hours to inhibit the growth of non-neuronal cells. The medium was replaced with fresh feeding medium 24 hours after adding AraC. After 3 DIV, half of the Neurobasal medium was replaced with fresh Neurobasal feeding medium every 4 days. 1 μl of the diluted AAV (4.8×10^12^ vg/ml) was dropped into the culture at 7 DIV and the degradation assay was performed at 13 or 14 DIV.

### Hippocampal acute slice preparation

Adult mice were anesthetized by isoflurane and fixed in a stereotaxic frame. The skull was exposed and a glass micropipette containing the AAV was inserted to dentate gyrus (AP = −2 mm; ML = +1.3 mm; DV = −2.05 mm). 500 nl of AAV was injected at 50 nl per min using a syringe pump (KD Scientific, Tokyo, Japan). 3 weeks after the AAV injection, a posterior brain block was cut into 300-μm thick coronal slices using a Vibratome VT1200S (Leica Microsystems, Wetzlar, Germany) in ice-cold oxygenated artificial cerebrospinal fluid (aCSF: 124 mM NaCl, 2.5mM KCl, 1.2 mM NaH_2_PO_4_, 24 mM NaHCO_3_, 5 mM HEPES, 13 mM glucose, 2 mM CaCl_2_. Slices were briefly transferred to an interface chamber containing oxygenated aCSF at RT. Slices were placed onto 35 mm glass base dish filled with Neurobasal feeding medium during the degradation assay.

### Immunocytochemistry

Cells were fixed in 4% paraformaldehyde in PBS for 10 min at RT. After fixation, cells were incubated in PBS containing 0.1% Triton X-100 for 15 min at RT. After washing twice with PBS, cells were blocked in a solution of PBS containing 5% normal donkey serum for 30 min at RT. Primary antibodies used are as follows: mouse anti-Tujl antibodies (Covance, 1:1,000); chicken anti-GFP antibodies (Abcam, 1:1,000). Fluorescent images of immunostained samples were obtained using a BZ-X700 microscope (Keyence, Osaka, Japan).

### Data acquisition and statistical analysis

Images were acquired using an FV1200 scanning confocal microscope (Olympus, Tokyo, Japan) equipped with diode lasers. For imaging primary culture, Z-series images (7 optical sections) were acquired with a 10 × water immersion objective lens (0.40 numerical aperture, Olympus). For imaging acute slice, Z-series images (5 optical sections) were acquired with a 20 × water immersion objective lens. GFP signal intensities within soma were measured with ImageJ (NIH, Bethesda, MD, USA). After subtracting background signals, signal intensity at each time point was normalized to the data at 10 min (100%). Statistical analyses were performed with OriginPro (OriginLab) software.

### IAA treatment

IAA (indole-3-acetic acid sodium salt, Sigma-Aldrich) was diluted in Neurobasal feeding medium to make 0.5 M stock solution. Pre-warmed IAA solution was applied in hippocampal neuronal cells and acute slice in a final concentration of 0.5 mM.

## Results

To test whether the AID system could be applicable in the nervous system, we initially used primary dissociated neurons. To avoid spontaneous basal degradation of degron-fused proteins suggested by a recent report [25], we cultured hippocampal neurons in culture media without serum. Two components, TIR1 from *Oryza sativa* (OsTIR1) and proteins fused with a mini-AID tag (mAID), were introduced into neurons by AAV viral vector infection. We generated AAV-DJ carrying *OsTIR1* and EGFP fused with *mAID* (pAAV-hSyn-OsTIR1-P2A-mAID-EGFP). Leucine-rich nuclear export signal (NES) sequence was also attached to the C-terminus region of EGFP to promote translocation of EGFP proteins into the cytoplasm. P2A peptide coding sequence was inserted into the middle of *OsTIR1* and *mAID-EGFP* to achieve simultaneous expression of these proteins (Fig. 1A). AAV-DJ containing solution (4.8 × 10^13^ vg/ml) was applied to dissociated hippocampal neurons prepared from PD0 pups (See Methods for details). Immunocytochemistry with antibodies against GFP and Tuj1 (a neuronal marker), revealed that mAID-EGFP proteins were present across the entire cytoplasm and that ectopic expression of mAID-EGFP proteins did not affect the morphology of the dissociated neurons (Fig. 1B).

**Figure 1.**
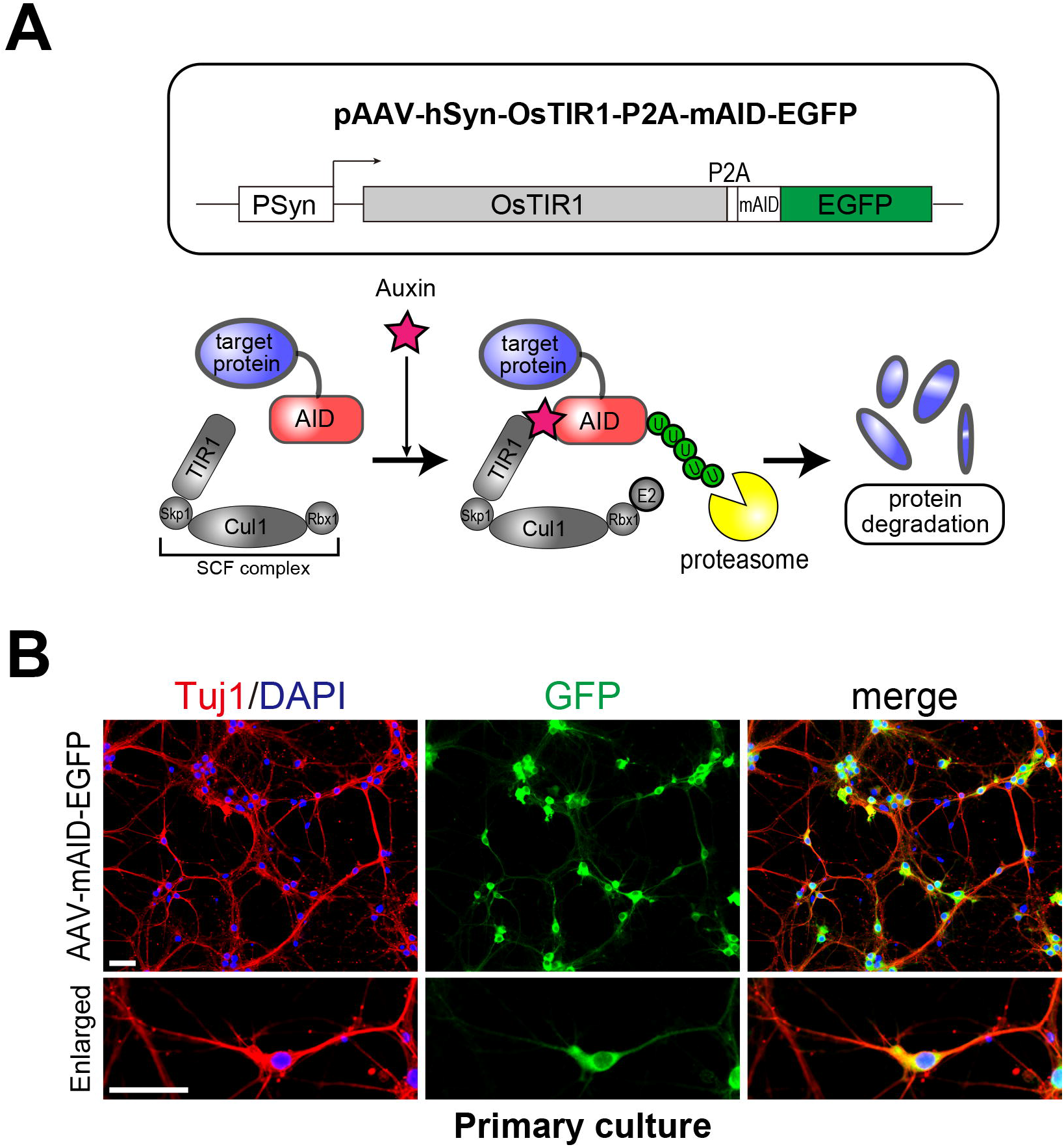
An auxin-inducible degron system for rapid protein depletion in neurons. **(A)** Schematic drawings of AAV vector for a protein degradation assay using the AID system (top). Virally expressed TIR1 proteins are incorporated into SCF complex (bottom left). In the presence of auxin, TIR1 interacts with AID-fused target proteins and promotes polyubiquitination by the E2 ligase (bottom middle). Ubiquitinated target proteins are rapidly degraded by the 26S proteasome (bottom right). **(B)** Transduction of dissociated hippocampal cells by the AID-fused EGFP. Hippocampal cultures transduced with AAV-hSyn-OsTIR1-P2A-mAID-EGFP vector were immunostained using antibodies against GFP and Tuj1 (a neuronal marker). Scale bars, 50 μm.

We then tested whether degradation of mAID-fused protein in dissociated neurons was triggered by application of indole-3-acetic acid (IAA). Dissociated hippocampal neurons prepared from PD0 pups were infected by the AAV-DJ carrying *OsTIRl* and *mAID-EGFP* at 7 days *in vitro* (7 DIV). One week after AAV infection (13-14 DIV), EGFP fluorescent intensities were quantified with fluorescence time-lapse imaging after IAA treatment to analyze the kinetics of protein degradation (Fig. 2A). We found that EGFP fluorescence dropped over time (Fig. 2B) and showed the weakest signal intensity 90 min after IAA application (Figs. 2C and D, n = 30 and 38 cells for EGFP and mAID-EGFP, respectively. \*\*\**p* < 0.001, Student’s *t*-test). In contrast, fluorescent signals from EGFP without mAID did not decrease upon IAA treatment. This result indicated that auxin-induced degradation of mAID-fused proteins occurred in *in vitro* dissociated hippocampal neurons.

**Figure 2.**
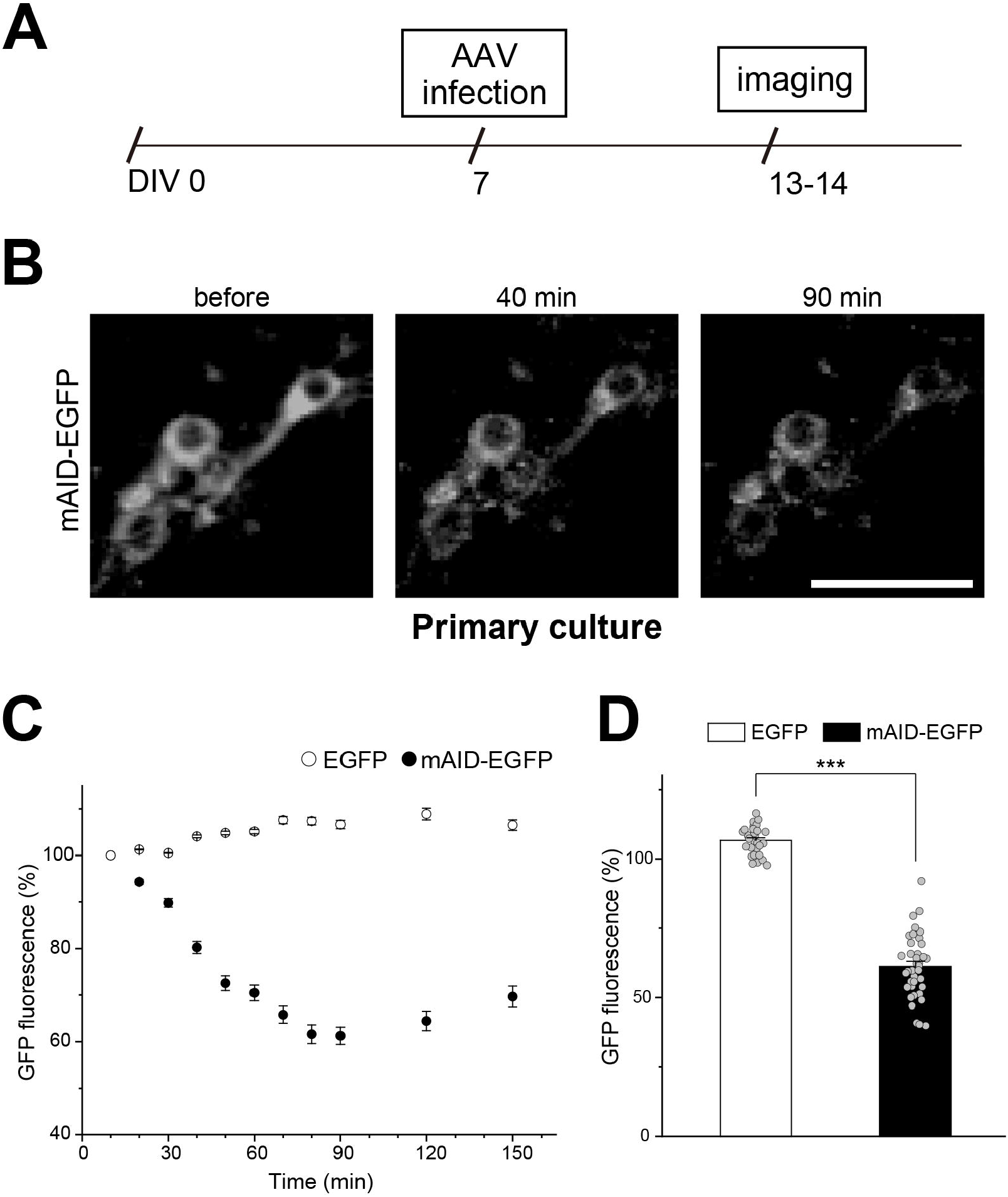
Auxin-induced protein degradation in dissociated hippocampal neurons. **(A)** Experimental paradigm for the protein degradation assay in dissociated neurons. Dissociated hippocampal neurons were infected with AAV-hSyn-OsTIR1-P2A-mAID-EGFP or AAV-hSyn-EGFP vectorat 7 DIV and the degradation assay was performed at 13 or 14 DIV. **(B)** Representative images of mAID-EGFP expressing dissociated hippocampal neurons before and after IAA treatment. GFP fluorescence decreased upon IAA treatment. Scale bar, 50 μm. **(C)** Signal intensities of GFP in hippocampal neurons were quantified and plotted with the min time point sample as 100%. Open circles indicate EGFP and filled circles indicate mAID-EGFP. **(D)** Signal intensities of EGFP in dissociated hippocampal neurons 90 min after IAA application. Error bars indicate the SEM. n = 30 and 38 cells for EGFP and mAID-EGFP, respectively. \*\*\**p* < 0.001, Student’s *t*-test.

We then tested whether the auxin-induced protein degradation occurs in the acute brain slices. We injected the AAV-OsTIR1-P2A-mAID-EGFP into the dentate gyrus of mice (Fig. 3A) and conducted protein degradation assay 3 weeks after AAV infection. Acute brain slices containing dentate gyrus were prepared and EGFP fluorescence signal intensities in hippocampal neurons with (+) or without (−) IAA treatment were measured by time-lapse imaging using confocal microscopy (Figs. 3B and C). We found that EGFP fluorescence decreased to about 50% of its initial value within 120 min after IAA application (Figs. 3C and D, n = 5 cells per group. \*\**p* < 0.01, Student’s *t*-test). This result implies that the AID system is partially working.

**Figure 3.**
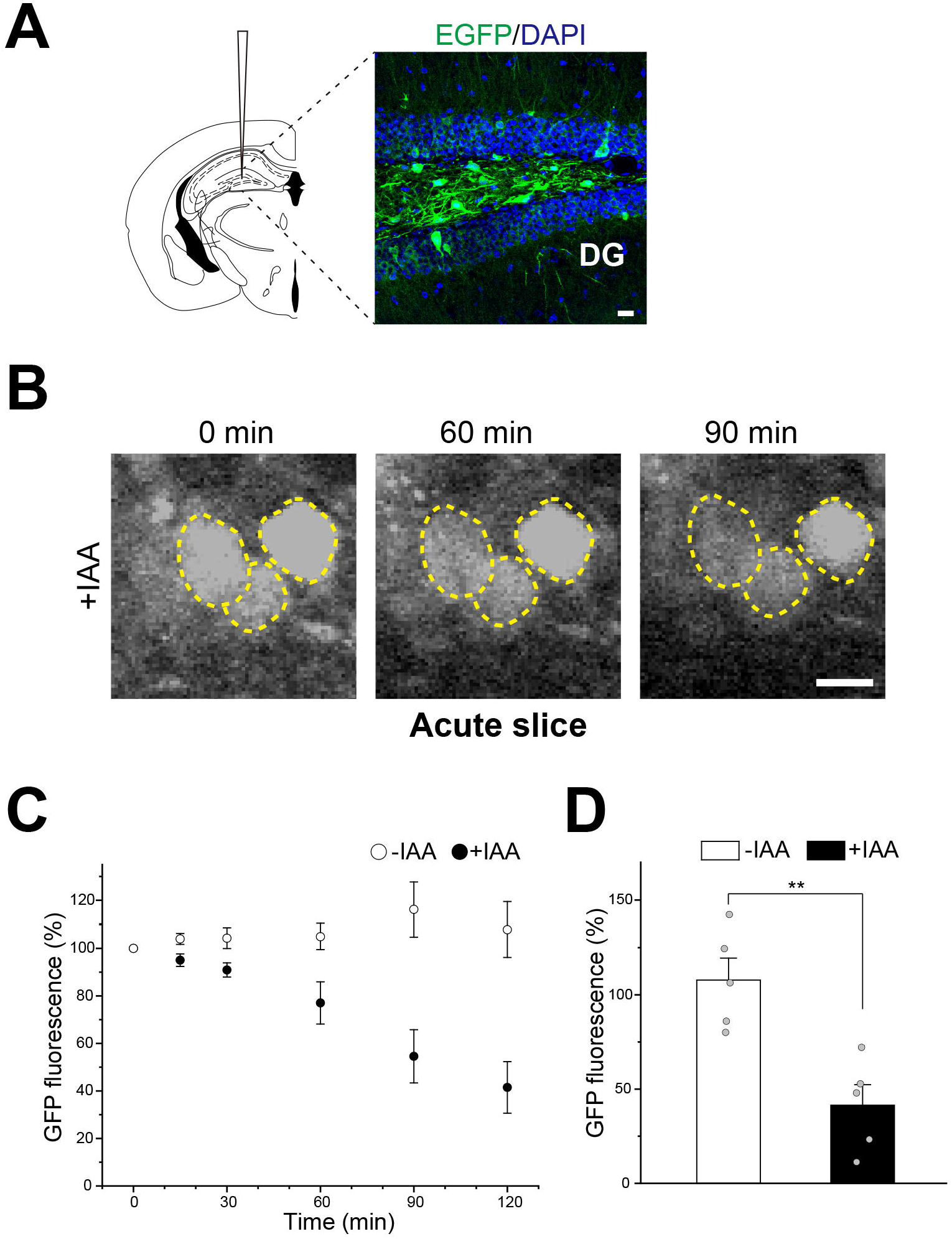
Auxin-induced protein degradation in acute brain slices. **(A)** AAV-hSyn-OsTIR1-P2A-mAID-EGFP vector was injected into dentate gyrus and the degradation assay was performed 3 weeks after the injection. Expression of EGFP in a coronal hippocampal section from the AAV-infected mouse. **S**cale bar, 20 μm. DG, dentate gyrus. **(B)** Representative images of mAID-EGFP expressing hippocampal neurons before and after IAA treatment. GFP fluorescence decreased upon IAA treatment. Neurons are demarked by dashed lines. **S**cale bar, 10 μm. **(C)** Signal intensities of mAID-EGFP in hippocampal neurons were quantified and plotted with the 0 min time point sample as 100%. **(D)** Signal intensities of GFP in hippocampal neurons 120 min after IAA application. Error bars indicate the SEM. n = 5 cells per group. \*\**p* < 0.01, Student’s *t*-test.

## Discussion

In the present study, we examined whether conditional protein degradation with the AID system can be transferable to the central nervous system. By time-lapse imaging of EGFP fluorescence, we concluded that auxin-inducible degradation of mAID-tagged proteins occurred in primary culture and acute slice conditions.

However, it should be noted that the efficiency and the rate of degradation were not comparable to those observed in other cell lines [18, 21]. Why is the AID system only partially working in neurons? One explanation is that the N-terminus fusion of mAID to EGFP was not suitable for efficient protein degradation. It might be possible that protein degradation occurs more efficiently and quickly if mAID was fused to the C-terminus region of the proteins. Another explanation is that the rate of protein synthesis was much higher than that of degradation induced by auxin. Because expression levels of virally-induced proteins are thought to be much higher than those of endogenous proteins, it is possible that virally-induced mAID-EGFP production overwhelmed protein degradation induced by auxin. It would be interesting to test the AID system combined with a knock-in approach to achieve physiological expression levels of mAID-fused proteins. Conditional and reversible control of specific proteins is desirable to elucidate brain functions. Further improvement in the AID system would provide powerful tools to investigate protein functions in the field of neuroscience.

## Acknowledgments

This work was supported by the Takeda Science Foundation, the Japan Foundation for Applied Enzymology, the Kato Memorial Bioscience Foundation, and the JSPS KAKENHI Grant Number 24370087, 15K21029, 18K14712. We appreciate Mr. Stuart Wax for his advice in the writing of this paper.

## Abbreviations

AID: auxin-inducible degron
EGFP: enhanced green fluorescent protein
SCF complex: Skp1-Cullin-F-Box protein complex
AAV: adeno-associated virus
DIV: day *in vitro*
IAA: indole-3-acetic acid sodium salt

## References

1. Capecchi MR: Gene targeting in mice: functional analysis of the mammalian genome for the twenty-first century. Nature reviews Genetics 2005, 6(6):507–512.

2. McManus MT, Sharp PA: Gene silencing in mammals by small interfering RNAs. Nature reviews Genetics 2002, 3(10):737–747.

3. Gaj T, Gersbach CA, Barbas CF, 3rd: ZFN, TALEN, and CRISPR/Cas-based methods for genome engineering. Trends Biotechnol 2013, 31(7):397–405.

4. Cong L, Ran FA, Cox D, Lin S, Barretto R, Habib N, Hsu PD, Wu X, Jiang W, Marraffini LA et al: Multiplex genome engineering using CRISPR/Cas systems. Science 2013, 339(6121):819–823.

5. Miller JC, Tan S, Qiao G, Barlow KA, Wang J, Xia DF, Meng X, Paschon DE, Leung E, Hinkley SJ et al: A TALE nuclease architecture for efficient genome editing. Nature biotechnology 2011, 29(2):143–148.

6. Urnov FD, Rebar EJ, Holmes MC, Zhang HS, Gregory PD: Genome editing with engineered zinc finger nucleases. Nature reviews Genetics 2010, 11 (9):636–646.

7. Hammond SM, Caudy AA, Hannon GJ: Post-transcriptional gene silencing by double-stranded RNA. Nature reviews Genetics 2001, 2(2): 110–119.

8. Banaszynski LA, Chen LC, Maynard-Smith LA, Ooi AGL, Wandless TJ: A rapid, reversible, and tunable method to regulate protein function in living cells using synthetic small molecules. Cell 2006, 126(5):995–1004.

9. Bonger KM, Chen LC, Liu CW, Wandless TJ: Small-molecule displacement of a cryptic degron causes conditional protein degradation. Nat Chem Biol 2011, 7(8):531–537.

10. Neklesa TK, Tae HS, Schneekloth AR, Stulberg MJ, Corson TW, Sundberg TB, Raina K, Holley SA, Crews CM: Small-molecule hydrophobic tagging-induced degradation of HaloTag fusion proteins. Nat Chem Biol 2011, 7(8):538–543.

11. Buckley DL, Raina K, Darricarrere N, Hines J, Gustafson JL, Smith IE, Miah AH, Harling JD, Crews CM: HaloPROTACS: Use of Small Molecule PROTACs to Induce Degradation of HaloTag Fusion Proteins. ACS Chem Biol 2015, 10(8): 1831–1837.

12. Nabet B, Roberts JM, Buckley DL, Paulk J, Dastjerdi S, Yang A, Leggett AL, Erb MA, Lawlor MA, Souza A et al: The dTAG system for immediate and target-specific protein degradation. Nat Chem Biol2018, 14(5):431–441.

13. Natsume T, Kanemaki MT: Conditional Degrons for Controlling Protein Expression at the Protein Level. Annu Rev Genet2017, 51:83–102.

14. Nishimura K, Fukagawa T, Takisawa H, Kakimoto T, Kanemaki M: An auxin-based degron system for the rapid depletion of proteins in nonplant cells. Nature methods 2009, 6(12):917–922.

15. Dharmasiri N, Dharmasiri S, Estelle M: The F-box protein TIR1 is an auxin receptor. Nature 2005, 435(7041):441–445.

16. Kepinski S, Leyser O: The Arabidopsis F-box protein TIR1 is an auxin receptor. Nature 2005, 435(7041):446–451.

17. Tan X, Calderon-Villalobos LI, Sharon M, Zheng C, Robinson CV, Estelle M, Zheng N: Mechanism of auxin perception by the TIR1 ubiquitin ligase. Nature 2007, 446(7136):640–645.

18. Holland AJ, Fachinetti D, Han JS, Cleveland DW: Inducible, reversible system for the rapid and complete degradation of proteins in mammalian cells. Proceedings of the National Academy of Sciences of the United States of America 2012, 109(49):E3350–3357.

19. Kanke M, Nishimura K, Kanemaki M, Kakimoto T, Takahashi TS, Nakagawa T, Masukata H: Auxin-inducible protein depletion system in fission yeast. BMC Cell Biol 2011, 12:8.

20. Zhang L, Ward JD, Cheng Z, Dernburg AF: The auxin-inducible degradation (AID) system enables versatile conditional protein depletion in C. elegans. Development 2015, 142(24):4374–4384.

21. Lambrus BG, Uetake Y, Clutario KM, Daggubati V, Snyder M, Sluder G, Holland AJ: p53 protects against genome instability following centriole duplication failure. The Journal of cell biology 2015, 210(1):63–77.

22. Iwasawa C, Narita M, Tamura H: Regional and temporal regulation and role of somatostatin receptor subtypes in the mouse brain following systemic kainate-induced acute seizures. Neuroscience research 2019.

23. Aurnhammer C, Haase M, Muether N, Hausl M, Rauschhuber C, Huber I, Nitschko H, Busch U, Sing A, Ehrhardt A et al: Universal real-time PCR for the detection and quantification of adeno-associated virus serotype 2-derived inverted terminal repeat sequences. Hum Gene Ther Methods 2012, 23(1):18–28.

24. Seibenhener ML, Wooten MW: Isolation and culture of hippocampal neurons from prenatal mice. J Vis Exp 2012(65).

25. Natsume T, Kiyomitsu T, Saga Y, Kanemaki MT: Rapid Protein Depletion in Human Cells by Auxin-Inducible Degron Tagging with Short Homology Donors. Cell Rep 2016, 15(1):210–218.

